# The Critical Period Microbiota Shape Brain Plasticity

**DOI:** 10.64898/2026.06.08.730811

**Authors:** Francesca Damiani, Kousha Changizi Ashtiani, Andrea Tognozzi, Sherif Abdelkarim, Sara Cornuti, Matteo Caldarelli, Pierre Baldi, Paola Tognini

**Affiliations:** Laboratory of Biology, Scuola Normale Superiore, Pisa, Italy; AI in Science Institute, School of Information and Computer Sciences, University of California, Irvine, CA, USA; Health Science Interdisciplinary Center, Scuola Superiore Sant’Anna, Pisa, Italy; Cardio-thoracic Department, Fondazione Toscana Gabriele Monasterio, Pisa, Italy

**Keywords:** Gut microbiota, Plasticity, Neurodevelopment, Critical Period, Visual Cortex, Antibiotic, Transcriptome, Blood-brain-barrier, Fecal Transplant

## Abstract

The gut microbiota is increasingly recognized as a regulator of brain function, yet its role in experience-dependent plasticity during postnatal development remains largely unknown. Here, we show that disrupting the gut microbiota with antibiotics during critical periods of visual cortex development impairs ocular dominance plasticity (ODP) in juvenile mice. Antibiotic treatment induces marked changes in microbial community composition and is accompanied by extensive transcriptional remodeling of the visual cortex, including pathways involved in extracellular matrix organization, blood-brain barrier function, and myelination. Remarkably, fecal transplantation of the juvenile microbiota into adult recipients restores ODP.

These findings identify the gut microbiota as a previously unrecognized regulator of neurodevelopmental plasticity and support the existence of microbiota-dependent critical periods of brain development. More broadly, our results suggest that early-life microbial perturbations may have lasting consequences for lifelong brain function and reveal that juvenile microbiota-derived signals could be exploited to promote plasticity in the adult brain.

## INTRODUCTION

Brain development is a prolonged and highly dynamic process which extends from gestation into early adulthood ^1^. While prenatal neurodevelopment is largely driven by genetic programs ^2–6^, postnatal brain maturation relies heavily on experience and on the heightened plasticity of neural circuits during critical periods (CP), when the complex interaction between environmental inputs and genetic factors exert maximal influence on circuit refinement ^1^. Disruption of these tightly regulated windows can lead to long-lasting alterations in brain function.

In parallel with postnatal brain development, the intestinal microbiota undergoes rapid maturation and remodeling during early life ^7,8^ ^9,10^. Gut microbial colonization begins at birth and is strongly shaped by maternal transmissions and early life exposure ^11,12^. The mode of delivery, feeding practices, and early environmental factors exposure ^13–17^ profoundly influence microbiota assembly, with long-term consequences for immune and metabolic health ^12,18–24^. Increasing evidence now suggests that this early microbial ecosystem may also contribute to postnatal brain development, positioning the gut microbiota as a potential regulator of experience-dependent neural plasticity.

During the neonatal period, the intestinal microbial ecosystem is unstable and of low diversity, progressively maturing with diet and environmental interactions ^25–27^. Breastfeeding promotes the colonization of beneficial taxa such as *Bifidobacterium*, whereas formula milk feeding leads to a more diverse but less specialized microbiota ^28–32^. Breastfeeding may be associated with improved neurodevelopment and cognitive outcomes ^33–37^. However, direct functional evidence linking early life microbiota integrity to specific neurodevelopmental processes remains limited. A major gut microbial ecological shift occurs at weaning, when the transition from milk to solid food reshapes the microbiome ^38–41^. This post-weaning period coincides with ongoing brain maturation and heightened plasticity and, therefore, potential vulnerability of neural circuits to environmental perturbations ^9,42^. The immaturity of the gut microbiota during post-weaning, infancy and childhood makes the community particularly susceptible to external insults, including antibiotic exposure. Antibiotics are widely prescribed in early life and are known to alter microbiota composition, diversity ^43–47^, and to alter cognitive and anxiety-like behaviour in mice ^48–51^. Yet the consequences of such perturbations on neural circuit functional plasticity during CPs remain poorly understood.

Here, we addressed this knowledge gap by investigating whether an intact intestinal microbiota is required for CP plasticity, using a well-established model: the rodent visual system. To mimic postnatal antibiotic exposure in infants, we treated juvenile mice with a broad-spectrum antibiotic cocktail (ABX) from weaning (P21) and for 10 days, corresponding to CP for ocular dominance plasticity (ODP). We found that ABX treatment impaired cortical plasticity and induced a profound transcriptional remodelling in the visual cortex. Genes associated with myelination, extracellular matrix (ECM) and basal membrane–related pathways were strongly impacted by ABX-driven microbial imbalance. Finally, to directly test the causal role of early-life microbiota in shaping cortical plasticity, we performed fecal microbiota transplantation (FT) from juvenile donors into adult, non-plastic recipient mice. Strikingly, transplantation of a “young” microbiota was sufficient to reinstate experience-dependent plasticity in the adult visual cortex.

## RESULTS

### One-week post-weaning ABX treatment dramatically affects fecal bacterial DNA content and gut microbiota composition

P21 mice were weaned and subjected to ABX dissolved in drinking water for 10 days, while control (CTRL) mice drank regular water (Fig. 1a).

**FIG.1.**
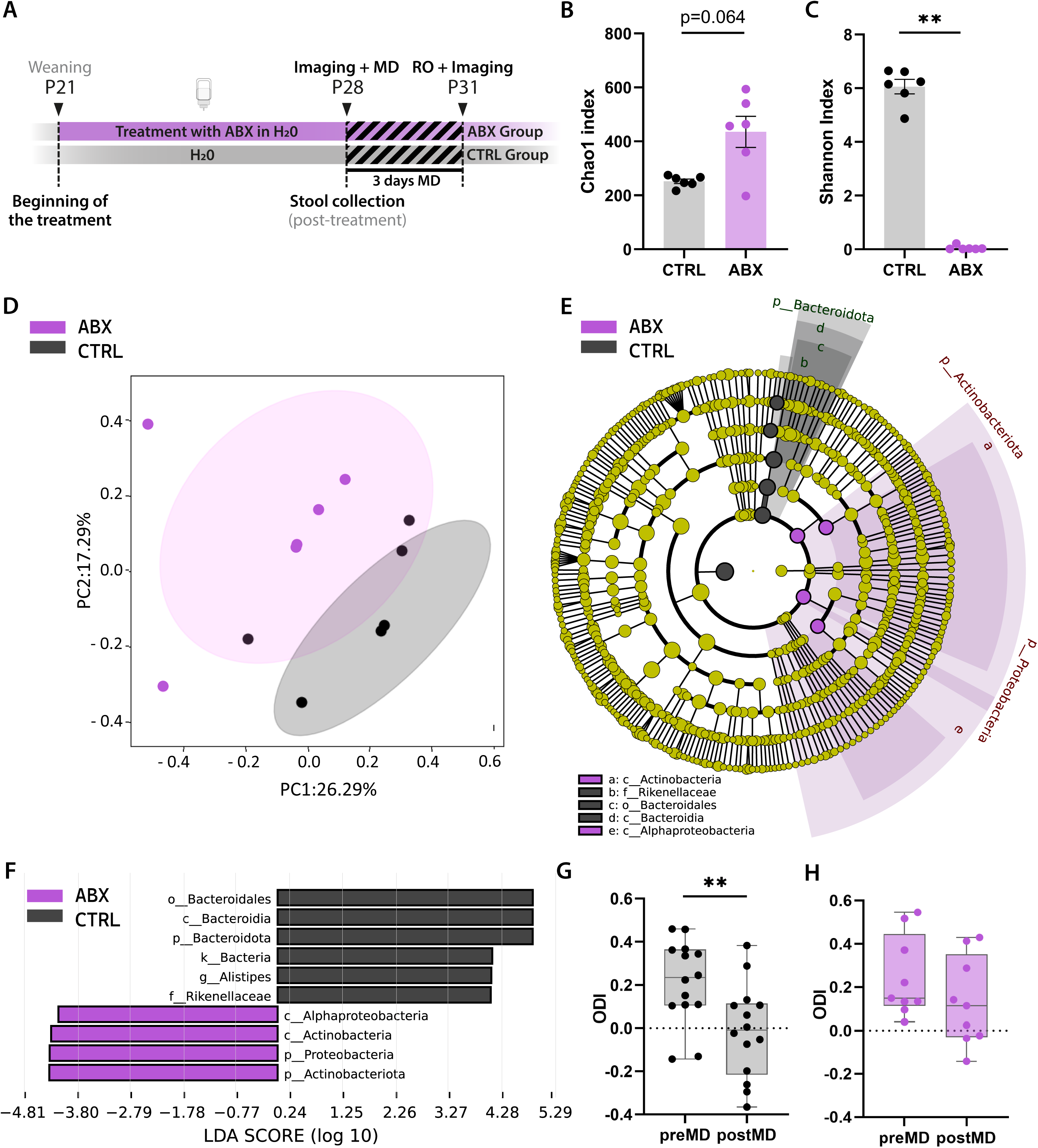
Post-weaning ABX administration reshapes gut microbiota composition and impairs ocular dominance plasticity during the critical period. **(A)** Timeline of the experiments. **(B-C)** Alpha diversity comparison between CTRL and ABX mice, expressed as **(B)** Chao1 and **(C)** Shannon indices. Chao1 index CTRL versus ABX Mann-Whitney U test p=0.064; Shannon index CTRL versus ABX Mann-Whitney U test **p=0.002. Error bars represent SEM. **(D)** PCoA plot showing beta diversity of the fecal microbiota from CTRL and ABX mice. The ellipses represent 95% confidence intervals for each group. Axes in the PCoA display the percentage of variation explained using Bray-Curtis dissimilarity. ANOSIM test: R=0.2, p=0.032. (**E-F**) LEfSe analysis of the fecal microbiota in CTRL and ABX mice. (**E**) Taxonomic cladograms resulted from LEfSe analysis of the gut microbiota, showing significantly differentially enriched taxa (relative abundance ≥ 0.5%) for taxonomy level L7 (species) between CTRL and ABX mice (**F**) Histogram of LDA scores (>2.5) obtained for bacterial taxa differentially abundant between CTRL and ABX mice. (G-H) Ocular Dominance Index (ODI) shifts after 3dMD in CTRL (**G**) and ABX (**H**) mice. ‘PreMD’ versus ‘postMD’ CTRL Paired t-test **p=0.006; ‘preMD’ versus ‘postMD’ ABX Paired t-test p=0.274. Whiskers indicate the minimum and maximum values; the median is displayed. Circles represent single experimental subjects (n=6-14 animals/group).

To assess the efficacy of ABX treatment, microbial DNA was extracted and quantified from fecal samples collected one week after the beginning of the treatment (P28, Fig. 1a). Analysis of fecal DNA revealed a significant reduction in DNA abundance in ABX-treated mice compared to CTRL mice drinking regular water (Suppl. Fig. 1a), confirming the successful manipulation of the gut microbiota. Additionally, ABX treatment caused a pronounced enlargement of the cecum, closely resembling the condition observed in GF mice and validating the effectiveness of ABX strategy in decreasing microbial abundance (Suppl. Figure 1b). Bacterial communities play a fundamental role in expanding the host’s metabolic capacity, and their depletion can result in reduced nutrient absorption and subsequent weight loss ^52^. Consistent with this evidence, ABX animals exhibited a slight, although significant reduction in body weight and a decreased body weight gain compared to CTRL mice at P28 (Suppl. Fig. 1c, d).

To investigate the composition of intestinal bacteria, fresh fecal samples were collected at P28 from both 1-week ABX-treated and CTRL mice, and 16S rRNA sequencing (seq) was performed. Alpha diversity, which represents the richness and evenness of species within a microbial ecosystem ^53^, was assessed using the Chao1 and Shannon indices. The Chao1 index, a measure of taxa richness, showed a trend toward a significant increase in alpha diversity in ABX-treated mice (Fig. 1b). In contrast, the Shannon index, which accounts for both taxa richness and evenness, was significantly lower in ABX mice compared to CTRL animals (Fig. 1c).

Beta diversity, reflecting the degree of phylogenetic similarity between microbial communities, was evaluated using the Bray-Curtis dissimilarity matrix. The principal coordinates analysis (PCoA) of the distance matrix displayed a distinct clustering of the ABX mice microbiota with respect to untreated CTRL (Fig. 1d).

To gain further insights into the effects of the ABX treatment on bacterial composition, we analyzed the relative abundance of the main bacterial families in CTRL and ABX mice. Across all samples, *Lachnospiraceae*, *Erysipelotrichaceae*, *Lactobacillaceae*, and *Muribaculaceae* were the dominant families, accounting for an average total abundance of 79.9% ± 14.4. Among these, the abundances of *Lachnospiraceae* and *Muribaculaceae* significantly declined in ABX-treated samples, while the abundance of *Erysipelotrichaceae* appeared to increase (Suppl. Fig. 1e). This shift is consistent with the observed decrease in Shannon index (Fig. 1c), indicating reduced evenness in the microbial community of ABX mice.

Finally, differences in the bacterial composition between ABX and CTRL mice were also analyzed by using the Linear discriminant analysis (LDA) of effect size (LEfSe) algorithms. LEfSe analysis revealed taxonomic dissimilarities between the two experimental groups. In particular, the class *Actinobacteria*, in the phylum Actinobacteriota, and the class *Alphaproteobacteria*, in the phylum Proteobacteria, were both enriched in the ABX-treated microbiota. On the other hand, the family Rikenellaceae, in the order Bacteroidales, was significantly decreased in the fecal microbiota of ABX mice compared to the CTRL condition (Fig. 1e,f).

In summary, ABX treatment led to substantial changes in the fecal bacterial composition, decreasing specific bacterial families (e.g. Lachnospiraceae, Muribaculaceae and Rikenellaceae families) while increasing other taxa, including Erysipelotrichaceae and Bacillaceae families, Actinobacteria and Alphaproteobacteria classes. Moreover, alpha-diversity data showed reduced taxa evenness, indicating that although overall diversity may have increased, the microbial community became more unevenly distributed.

### Post-weaning microbial manipulation impairs ocular dominance plasticity during the critical period

After validating the efficacy of our ABX administration, we tested whether an intact gut microbiota is necessary for experience-dependent plasticity in the juvenile visual cortex. Following one week of ABX administration, we assessed ODP by IOS imaging. In particular, visual cortical responses were analysed before and after 3 days of monocular deprivation (MD) in mice at P28-P31 (Fig. 1a). Since ODP in mice typically peaks between P28 and P31, a significant shift in the ocular dominance index (ODI) towards the open eye (ipsilateral to the recorded visual cortex) was anticipated in the CTRL group (Fig. 1g). Strikingly, ABX-treated mice did not display OD shift (Fig. 1h), indicating an impairment in ODP mechanisms.

### ABX-driven microbial manipulation reprograms the transcriptome in the mouse visual cortex

To explore the potential mechanisms underlying the effect of microbiota manipulation on CP plasticity, we performed an RNA sequencing (RNA-seq) in the visual cortex of ABX and CTRL mice.

Strikingly, transcriptome analysis revealed a total of 1083 genes to be differentially expressed between CTRL and ABX mice. In particular 443 transcripts were upregulated and 640 transcripts were downregulated in the visual cortex of the ABX-treated animals compared with the CTRL group (Fig. 2a, b). For the complete list of transcripts see Suppl. Table 1.

**FIG.2.**
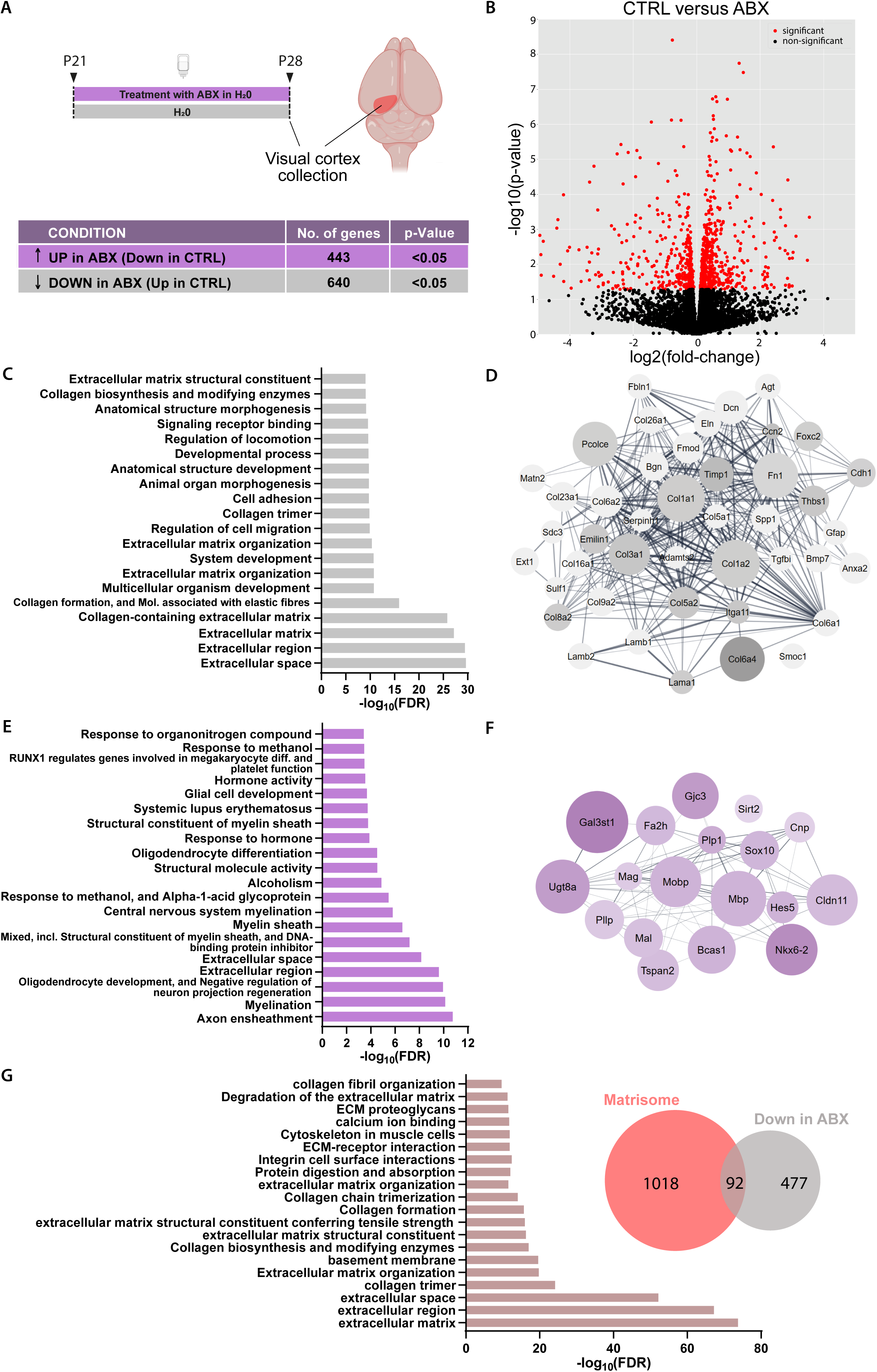
ABX-driven microbiota manipulation induces profound transcriptional changes in the visual cortex of critical period mice. (A) Schematic illustration of the experiment (top) and table summarizing genes significantly upregulated (purple) or downregulated (gray) in the visual cortex following ABX treatment. Cyber t-test p<0.05. N=4 animals/group. **(B)** Volcano plot showing upregulated (left part) and downregulated (right part) genes in the ABX group with respect to CTRL. **(C-F)** GO analysis on differentially expressed genes between CTRL and ABX. **(C)** Top 20 GO annotations related to downregulated genes in the ABX group. **(D)** Reactome pathway of genes downregulated in the ABX group and belonging to extracellular matrix terms. Colors: Log2(fold-change). Circle dimensions: Log10(p-value). **(E)** Top 20 GO annotations related to upregulated genes in the ABX group. **(F)** Reactome pathway of genes upregulated in the ABX group and belonging to myelination terms. Colors: Log2(fold-change). Circle dimensions: Log10(p-value). **(G)** Top 20 GO annotations for genes overlapping between the matrisome and downregulated gene set in the ABX group (left), and corresponding Venn diagram displaying the overlap (right).

To gain further insight on the biological processes impacted by microbiota manipulation, we performed a gene ontology (GO), Kyoto Encyclopedia of Genes and Genomes (KEGG) and Reactome pathway enrichment analysis using the DAVID algorithms. Enrichment analysis of transcripts downregulated in the ABX-treated mice revealed multiple annotations linked to the ECM, such as “*Extracellular matrix organization*”, “*Collagen formation*”, “*Collagen-containing Extracellular matrix*”, “*Extracellular matrix space*” and “*Extracellular matrix region*” (Fig. 2c, d and suppl. Table 2). Moreover, the same analysis performed on transcripts upregulated in the ABX visual cortex unveiled a strong overrepresentation of pathways involved in myelination, including “*Structural constituent of myelin sheath*”, “*Oligodendrocyte differentiation*”, “*Central nervous system myelination*”, “*Myelin sheath*”, “*Structural constituent of myelin sheath*” and “*Oligodendrocyte development*” (Fig. 2e, f and Suppl. Table 2). Notably, both myelination and ECM remodelling have been previously involved in experience-dependent plasticity^54–62^. Since studies in the mammalian visual cortex have shown that increased myelination and ECM remodeling limit CP plasticity ^63^, our findings suggest that the gut microbiota during the CP may play a role in regulating ODP by influencing these processes.

To further characterize the transcriptomic signature related to ECM, we crossed the ABX downregulated transcripts with the matrisome list ^64^. Out of 640 downregulated transcripts in ABX condition, 92 genes overlapped with the matrisome (Fig. 2g). Notably, Functional annotation enrichment analysis revealed a significant overrepresentation of extracellular-related categories, including multiple collagen and ECM related annotations, “*integrin cell surface interaction*” and “*basement membrane*” (Fig. 2g and Suppl. Table 2). As the basement membrane is a non-cellular component of the blood brain barrier (BBB), which contributes to its stability and regulates its permeability ^65^, these data may point toward alterations in BBB integrity following ABX-induced microbiota remodelling during the CP.

### ABX treatment affects ECM organization, BBB permeability and myelination pathways

Based on the observed transcriptome reprogramming indicating potential alterations in the ECM of ABX mice, we first investigated the visual cortex PNN as they are crucial extracellular structures involved in experience-dependent plasticity ^60^. Since PNN assembly has been linked to the maturation of parvalbuminergic (PV^+^) neuronal network, which is involved in restricting ODP ^66,67^, we simultaneously analysed PV^+^ interneurons distribution in the visual cortex (Fig. 3a).

**FIG.3.**
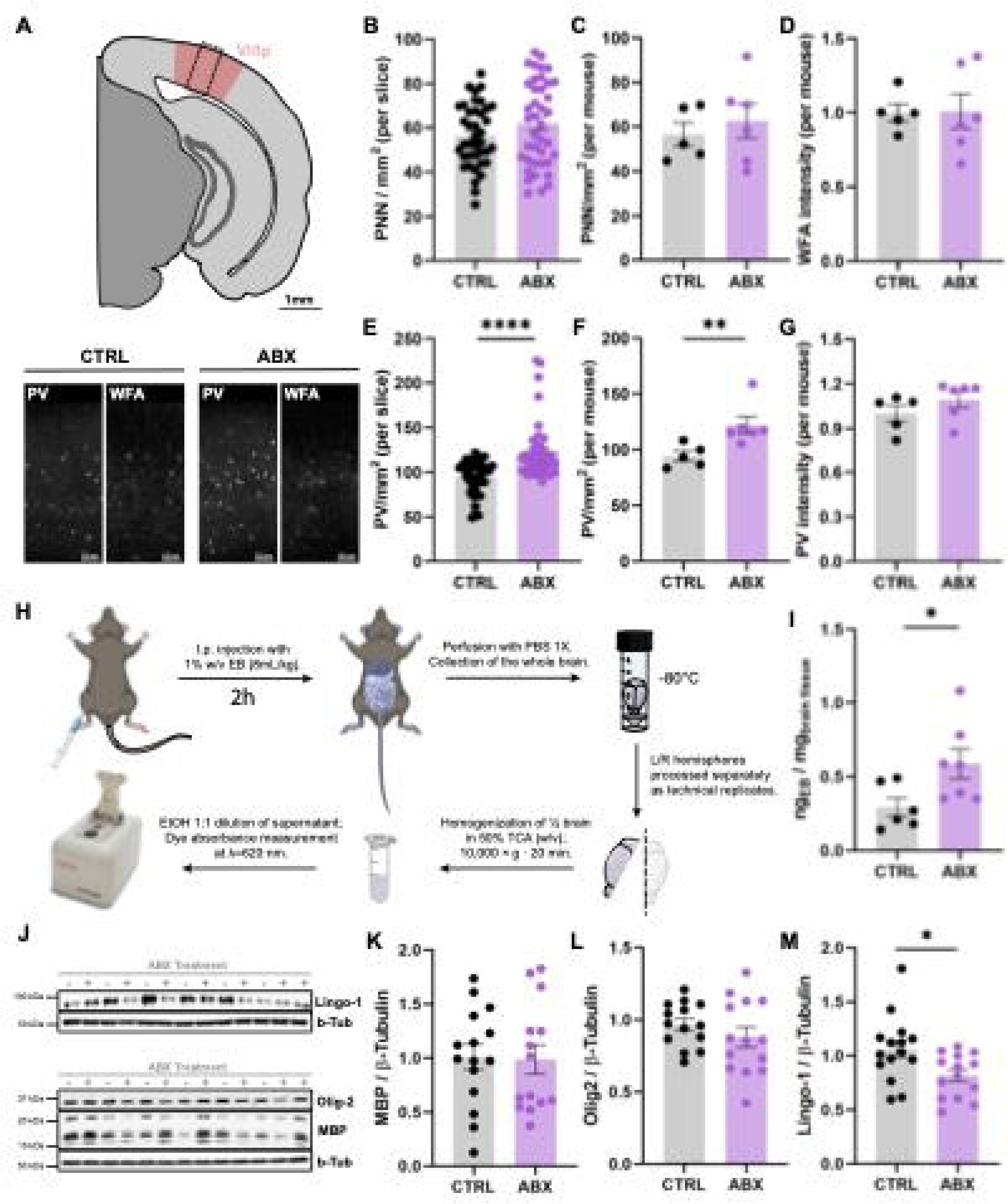
Antibiotic cocktail treatment affects PV^+^ interneurons density, blood-brain barrier integrity and cortical myelination-related proteins in the juvenile visual cortex. **(A-G)** Immunofluorescence (IF) analysis on the V1 region in CTRL and ABX mice stained for PV and WFA. N=8-10 slices/animal, corresponding to N=5-6 animals/group. **(A)** Representative images of PV and WFA staining in the V1 region of CTRL (bottom left) and ABX (bottom right) mice. **(B-D)** Quantitative IF analysis of WFA^+^ perineuronal nets (PNN). **(B)** WFA^+^ PNN density (PNN/mm2) per slice. Mann-Whitney U test, p=0.219. **(C)** Average WFA^+^ PNN density (PNN/mm2) per mouse. Unpaired t-test, p=0.552. **(D)** Average WFA staining intensity per mouse. Unpaired t-test, p=0.953. **(E-G)** Quantitative IF analysis of PV^+^ interneurons. **(E)** PV^+^ interneurons density (PV/mm2) per slice. Mann-Whitney U test, ****p<0.0001. **(F)** Average PV^+^ interneurons density (PV/mm2) per mouse. Mann-Whitney U test, **p=0.008. **(G)** Average PV staining intensity per mouse. Mann-Whitney U test, p=0.177. **(H,I)** Evans Blue (EB) dye permeability assay. N=6/7 animals/group. **(H)** Schematic representation of the experimental protocol. **(I)** Quantification of EB extravasation in the brains of CTRL and ABX mice, expressed as μg dye/mg brain tissue. Unpaired t-test, *p=0.036. **(J-M)** Western blot analysis of MBP, Olig2, and Lingo-1 protein expression in visual cortex extracts from CTRL and ABX mice. β-tubulin was used as a loading control. N=14–15 animals/group. **(J)** Representative western blot images. **(K–M)** Densitometric quantification of protein expression, shown as the ratio of band intensity for **(K)** MBP/β-tubulin, **(L)** Olig2/β-tubulin, and **(M)** Lingo-1/β-tubulin. MBP/β-tubulin ‘CTRL’ versus ‘ABX’ Unpaired t-test p=0.891; Olig2/β-tubulin ‘CTRL’ versus ‘ABX’ Unpaired t-test p=0.286; Lingo1/β-tubulin ‘CTRL’ versus ‘ABX’ Unpaired t-test *p=0.018. Error bars represent SEM. Circles represent single experimental subjects, unless otherwise specified.

No differences were observed in PNN density or intensity (Fig. 3b-d). On the other hand, the ABX mice visual cortex displayed an increase in PV+ neuron density, but not intensity, compared to the CTRL group (Fig. 3e-g). This data suggests a potential dysregulation in postnatal development of inhibitory circuits upon ABX-driven intestinal microbiota manipulation.

Then, we focused on BBB integrity. Evaluation of Evans blue dye extravasation revealed a significant increase in BBB permeability in ABX treated mice compared with CTRL mice (Fig. 3h, i).

Because the RNA-req analysis showed a significant overrepresentation of myelin-related annotations in the visual cortex of ABX mice, suggesting altered myelination, we further examined the levels of key myelination- related proteins. We analysed two positive regulators of myelination, Oligodendrocyte transcription factor 2 (Olig2) ^68–70^, and myelin basic protein (MBP) ^71–73^, as well as the negative regulator of oligodendrocyte differentiation Leucine rich repeat and ig domain containing 1 (Lingo1) protein ^74^. While MBP and Olig2 were not altered (Fig 3j-l), we found a significant reduction in visual cortical Lingo1 protein in ABX condition with respect to the CTRL group (Fig. 3j, m).

Overall, these findings demonstrate that an intact gut microbiota is essential for cortical plasticity and critical period regulation and suggest that ABX-induced dysbiosis may impact key neurodevelopmental processes, such as PV network and myelin maturation.

### Critical period mice exhibit a distinct gut microbiota composition compared to adulthood

To further test the hypothesis that the “young” gut microbiota could mediate ODP during the CP, we first analysed whether the CP intestinal microbiota composition differed from that of adult mice, which do not exhibit ODP. The analysis of alpha diversity revealed a significantly lower Chao1 index in young compared to adult mice, while the Shannon index was not different (Fig. 4a, b). This finding likely reflects the ongoing maturation and growing complexity of the young gut microbiota during the post-weaning period, when the bacterial ecosystem is characterized by fewer bacterial taxa ^39^. Moreover, beta diversity analysis displayed distinct clustering between young and adult mice (Fig. 4c), suggesting a different phylogenetic signature.

**FIG.4.**
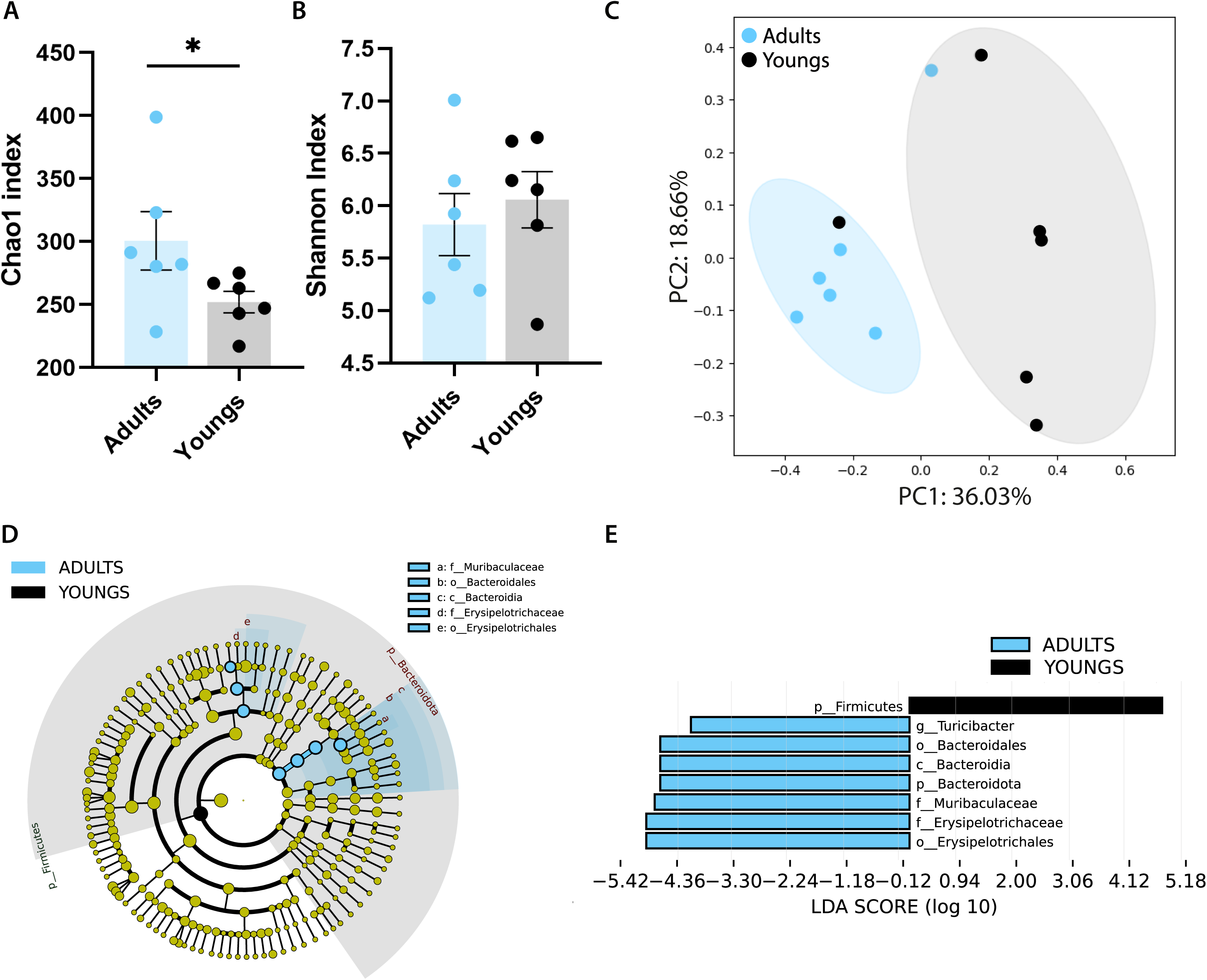
Characterization of the gut microbiota in young and adult mice reveals significant differences between the two ages. **(A,B)** Alpha diversity comparison between young and adult mice, expressed as **(A)** Chao1 and **(B)** Shannon indices. Chao1 index ‘Adults’ versus ‘Youngs’ Mann-Whitney U test *p=0.041; Shannon index ‘Adults’ versus ‘Youngs’ Mann-Whitney U test p=0.588. **(C)** PCoA plot showing beta diversity of the fecal microbiota of young and adult mice. The ellipses represent 95% confidence intervals for each group. Axes in the PCoA display the percentage of variation explained using Bray-Curtis dissimilarity. ANOSIM test: R=0.485, p=0.012. (**D-E**) LEfSe analysis of the fecal microbiota in young and adult mice. (**D**) Taxonomic cladogram showing significantly differentially enriched taxa (relative abundance ≥ 0.5%) for taxonomy level L7 (species) in young and adult mice. (**E**) Histogram of LDA scores (>2.5) obtained for bacterial taxa differentially abundant in juvenile and adult animals. Error bars represent SEM; circles represent single experimental subjects (n=6 animals/group).

Investigation on the relative abundance of major bacterial families revealed that Lactobacillaceae and Lachnospiraceae were enriched in young mice, whereas Erysipelotrichaceae and Muribaculaceae were more abundant in the adult group (Suppl Fig.2).

Consistent with these findings, LEfSe analysis identified strong differences between the young and adult microbiota composition (Fig. 4d,e). Specifically, the phylum *Firmicutes* was significantly enriched in young animals, while the genus *Turicibacter*, in the family Turicibacteraceae, in the order Erysipelotrichales, the family Erysipelotrichaceae, in the order Erysipelotrichales, and the family Muribaculaceae, in the order Bacteroidales, were more abundant in adults (Fig. 4 d,e).

Overall, these results confirm our previous findings ^75^ and several studies in both humans and mouse models ^76–78^, highlighting substantial differences in gut microbiota composition between young and adult mice.

### The “critical period” gut microbiota is sufficient to promote neuroplasticity in adult mice

After receiving proof of distinct microbiota composition in young and adult animals, and in order to find a causal link between CP plasticity and intestinal microbe features, we conducted a fecal microbiota transplantation (FT) experiment. Adult recipient mice (4 months old) received feces from young donor mice being at the peak of CP (∼P28-30) (FT-CP). Control groups consisted of adult recipients that either received a FT from adult conspecific mice (FT-AD) or were treated with the transplantation vehicle (FT-PBS). To assess whether the “young” microbiota could reopen the CP for ODP in adults, recipient mice underwent IOS imaging before and after three days of MD (Fig. 5a). Strikingly, while as expected no significant differences were present in the post-MD ODI of both control groups, FT-AD and FT-PBS (Fig. 5b, c), a significant OD shift was observed in the mice receiving a “critical period” gut microbiota (Fig. 5d).

**FIG.5.**
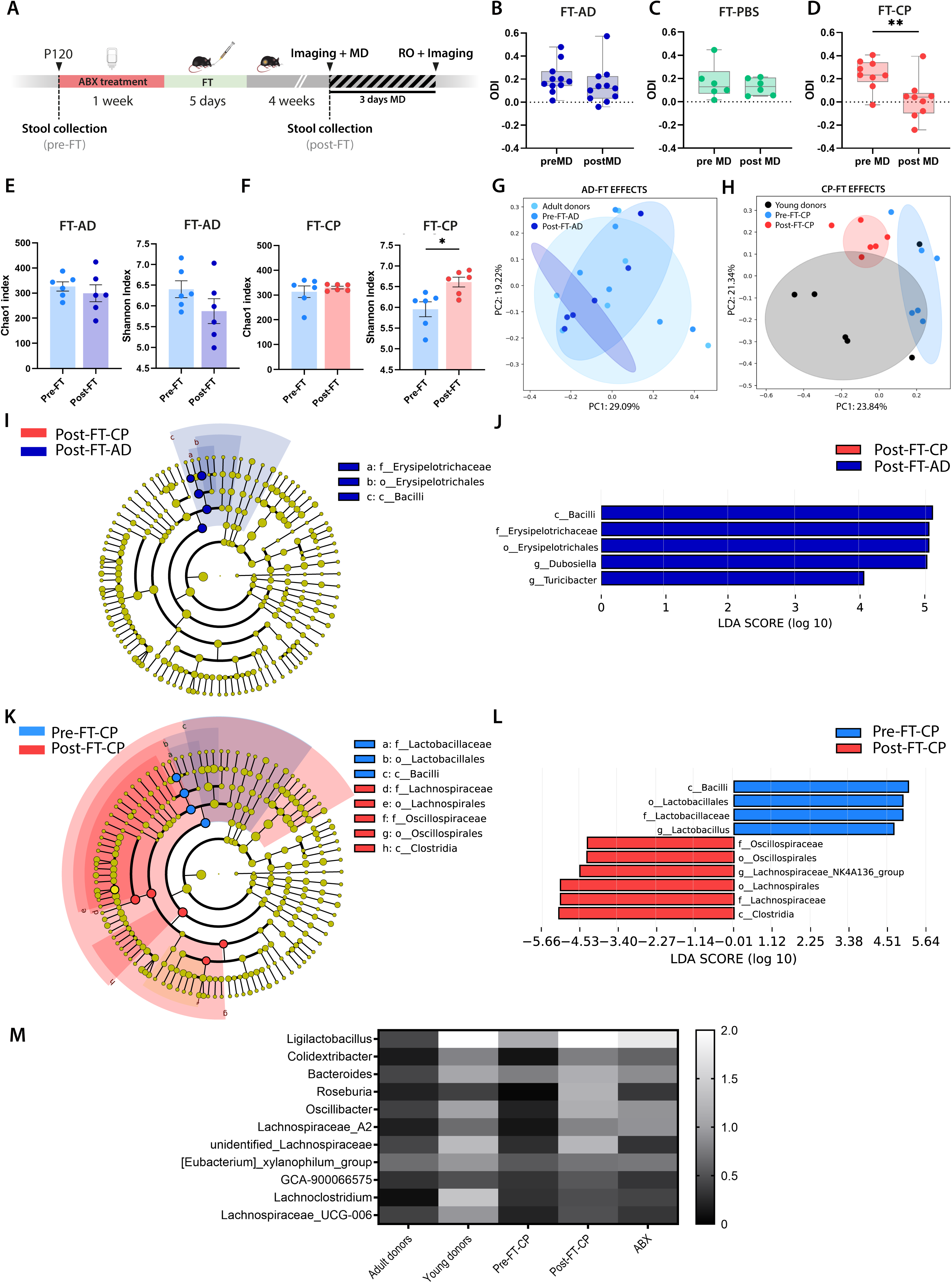
Fecal microbiota transplant from juvenile donors restores cortical plasticity in adult recipients and successfully colonizes the adult intestine. **(A)** Experimental timeline. **(B-D)** OD shifts after 3dMD in adult recipient mice of **(B)** adult donors (FT-AD), **(C)** PBS vehicle (FT-PBS), and **(D)** young donors. ‘PreMD’ versus ‘postMD’ FT-AD Wilcoxon matched-pairs signed rank test p=0.147; ‘preMD’ versus ‘postMD’ FT-PBS Paired t-test p=0.518; ‘preMD’ versus ‘postMD’ FT-CP Paired t-test **p=0.007. **(E,F)** Alpha diversity comparison between ‘Pre-FT’ and ‘Post-FT’ in **(E)** FT-AD and **(F)** FT-CP mice, expressed as Chao1 and Shannon indices. FT-AD Chao1 index ‘pre-FT’ versus ‘post-FT’ Wilcoxon matched pairs signed rank test p=0.843; FT-AD Shannon index ‘pre-FT’ versus ‘post-FT’ Wilcoxon matched pairs signed rank test p=0.218. FT-CP Chao1 index ‘pre-FT’ versus ‘post-FT’ Wilcoxon matched pairs signed rank test p=0.843; FT-CP Shannon index ‘pre-FT’ versus ‘post-FT’ FT-CP Wilcoxon matched pairs signed rank test *p=0.031. **(G, H)** PCoA plots showing beta diversity between donors and their corresponding pre- and post-FT recipient mice. The ellipses represent 95% confidence intervals for each group. Axes in the PCoA display the percentage of variation explained using Bray-Curtis dissimilarity. **(G)** PcoA on the ‘AD-FT effects’ showing overlap between adult donors and their recipient mice pre- and post-FT. ANOSIM test: ‘Adult donors’ versus ‘Pre-FT-AD’ R=-0.074, p=0.695; ‘Adult donors’ versus ‘Post-FT-AD’ R=-0.042, p=0.674; ‘Pre-FT-AD’ versus ‘Post-FT-AD’ R=0.072, p=0.211. **(H)** PCoA on the ‘CP-FT effects’ showing distinct clusters between young donors and their recipient mice pre-FT and a partial overlap with their recipient mice post-FT. ANOSIM test: ‘Young donors’ versus ‘Pre-FT-AD’ R=0.361, p=0.031; ‘Young donors’ versus ‘Post-FT-AD’ R=-0.485, p=008; ‘Pre-FT-AD’ versus ‘Post-FT-AD’ R=0.518, p=0.001. **(I-L)** LEfSe analysis of the fecal microbiota in FT recipient mice. (**I**) Taxonomic cladogram showing significantly differentially enriched taxa (relative abundance ≥ 0.5%) for taxonomy level L7 (species) between post-FTs of recipient mice receiving FT from either young or adult donors. (**J**) Histogram of LDA scores (>2.5) obtained for bacterial taxa differentially abundant between post-FTs of recipient mice receiving FT from either young or adult donors. (**K**) Taxonomic cladogram showing significantly differentially enriched taxa (relative abundance ≥ 0.5%) for taxonomy level L7 (species) between pre-FT and post-FT from young donors. (**L**) Histogram of LDA scores (>2.5) obtained for bacterial taxa differentially abundant between pre-FT and post-FT from young donors. (**M**) Heatmap displaying the relative abundance of the 11 potential “pro-plasticity” bacterial taxa (genus level) across all experimental groups. Each row represents a bacterial taxon, and each column corresponds to an individual experimental group. Color intensity indicates the relative abundance expressed as percentage, with darker shades representing lower abundance and lighter shades indicating higher abundance (white=relative abundance≥2%). Error bars represent SEM; circles represent single experimental subjects (n=6/10 animals/group).

This result demonstrates that transferring the young intestinal microbiota is sufficient to restore experience-dependent plasticity in the adult visual cortex. Also, this data reinforces the concept that an intact intestinal microbiota during CP is necessary for a correct establishment of experience-driven plasticity in postnatal neural circuits.

### Fecal microbiota transplant enables effective transfer of key ‘critical period’ gut microbial taxa

To identify the pro-plasticity gut microbial signature in the FT experiment, fresh fecal samples were collected from recipient mice, before and after the FT, and 16S rRNA-seq profiles were compared within recipients and with the donor groups, corresponding to adult and young mice whose microbiota is described in Fig. 4.

No significant differences were found in either the Chao1 and Shannon index when comparing pre-FT and post-FT samples from FT-AD mice (Fig. 5e), nor in the Chao1 index for pre-FT and post-FT samples from FT-CP mice (Fig. 5f). However, the Shannon index showed a significant increase in the post-FT samples from FT-CP mice (Fig. 5f). Beta diversity between the gut microbiota of adult donors and corresponding recipient mice, before and after FT, exhibited similar clustering patterns (Fig. 5g). On the other hand, the PCoA plot of the fecal microbiota from young donors and their corresponding recipient mice displayed diverse clustering when compared to the pre-FT microbiota (Fig.5h). Importantly, after FT, the microbiota of FT-CP partially overlapped with the microbiota of their young donors and differentiated from the pre-FT condition (Fig. 5h). Relative abundance analysis identified Lachnospiraceae, Erysipelotrichaceae, Lactobacillaceae, and Muribaculaceae as the predominant microbial families across all samples, with an average total abundance of 88.0% ± 10.5 (Suppl. Fig. 3). Notably, FT-CP exhibited a distinct shift in microbial composition, characterized by an increase in Lachnospiraceae and a decrease in Erysipelotrichaceae, closely resembling the microbiota profile of the young donors (Supp. Fig. 2). LEfSe analysis of post-FT recipient mice revealed that the Erysipelotrichaceae family, which was reduced in young donors compared to adult donors, was also decreased in post-FT-CP mice relative to post-FT-AD mice (Fig. 5i, j). This finding supports the successful transfer of microbial communities. Importantly, LEfSe analysis revealed significant differences in the gut microbiota composition of pre- and post-FT-CP recipient mice (Fig. 5k, l). In particular, post-FT-CP mice displayed an increase in the family Lachnospiraceae, in the order Clostridiales, and in the family Oscillospiraceae, in the order of Oscillospirales, both within the class of Clostridia, alongside a reduction in the abundance of the family Lactobacillaceae, in the order Lactobacillales, in the class Bacilli (Fig 5k, j). Importantly, Lachnospiraceae and Oscillospiraceae were also enriched in young donors compared to the adults.

Finally, we identified potential “pro-plasticity” bacterial taxa by comparing their relative abundance across adult donors, young donors, pre-FT-CP, post-FT-CP and ABX-treated mice. Specifically, we performed the analysis at the genus level, using as a selection criterion those taxa that were more abundant in young donors and post-FT-CP, compared to adult donors, pre-FT-CP and ABX-treated groups. Eleven bacterial taxa were identified: *Ligilactobacillus, Colidextribacter, Bacteroides, Roseburia, Oscillibacter, Lachnospiraceae_A2, unidentified_Lachnospiraceae, [Eubacterium]_xylanophilum_group, GCA-900066575, Lachnoclostridium and Lachnospiraceae_UCG-006* (Fig. 5m). Notably, *Ligilactobacillus, Bacteroides, Roseburia, Oscillibacter, [Eubacterium] xylanophilum group, Lachnoclostridium* and *Lachnospiraceae_UCG-006* are all genus associated with SCFA production ^79–88^.

Altogether, 16S rRNA-seq data demonstrated the successful engraftment of the gut microbiota from young donors into FT-CP recipient mice, reinforcing a causal link between intestinal microbe composition and CP plasticity. Importantly, the identification of 11 bacterial taxa enriched in young donors and post-FT-CP mice suggests a potential microbial signature associated with enhanced plasticity.

## DISCUSSION

Growing evidence suggests that the gut microbiota may play a role in neuroplasticity through multiple mechanisms, including epigenetic regulation, neurotransmitter synthesis and microglial modulations ^9^. However, its specific influence on experience-dependent plasticity during sensitive windows of postnatal neural network maturation, remains largely unexplored. Here, we show for the first time that ABX-treatment during CP impairs functional plasticity phenomena potentially through the modulation of myelination, BBB permeability and inhibitory circuit maturation. Importantly, we demonstrate that the postweaning intestinal microbiota is not only necessary and crucial for CP plasticity but it is also sufficient to promote plasticity in adulthood, “rejuvenating” the neural circuits. Taken together, our work suggests a novel concept: the existence of “microbiota-dependent critical periods”.

### Antibiotic-driven alterations in gut microbiota composition impact experience-dependent plasticity and reshape the visual cortex transcriptome

Antibiotics represent one of the greatest advancements in modern medicine, revolutionizing the treatment of bacterial infections and significantly reducing mortality ^89^. However, their excessive use, particularly in infants during the perinatal and postnatal period, has been linked to changes in the maturation trajectory of the intestinal microbiota, with potentially long-term consequences on immune system function and metabolic regulation ^90–93^. To mimic early-life antibiotic exposure in infants, mice received ABX immediately after weaning for the subsequent 10 days. In rodent models, this time-window represents the CP for ODP ^94,95^, and coincides with the transition from maternal milk to solid food, a key developmental interval for shaping the intestinal microbiota ^7^.

As expected, ABX treatment profoundly altered the gut microbiota composition, leading to a significant increase in taxa richness but a decrease in their evenness, suggesting that dominant commensal bacteria were suppressed, potentially allowing the expansion of opportunistic or antibiotic-resistant species. Principal coordinates analysis confirmed a distinct clustering of ABX mice compared to CTRL, reflecting substantial shifts in microbial composition. Notably, ABX administration reduced the relative abundance of beneficial bacterial families such as Lachnospiraceae, Muribaculaceae and Rikenellaceae, which are involved in fiber fermentation and the production of SCFA ^83,96–99^ with neuroprotective and anti-inflammatory properties ^100–103^. Conversely, we observed an expansion of potentially detrimental taxa, including Erysipelotrichaceae, which have been linked to inflammatory conditions and metabolic disorders ^104–106^. Additionally, we detected an increase in Alphaproteobacteria, a group with an unclear role in intestinal homeostasis but also associated with gut dysbiosis ^106,107^. These findings indicate that ABX administration induced a profound microbial composition reprogramming, shifting the gut ecosystem toward a state characterized by taxa linked to inflammation and dysbiosis. This shift may significantly affect host physiology, leading to long-term consequences, including altered brain maturation and experience-dependent plasticity. To test this hypothesis, we assessed ODP at the peak of the CP. ABX treated mice did not exhibit the OD shift supposed to be present at that age, indicating an impairment in visual cortical plasticity. Thus, an intact gut microbiota is necessary for experience-dependent plasticity in the juvenile brain, a fundamental process for brain postnatal development. Importantly, this piece of data could be relevant to inform the clinic over the excessive use of antibiotics in neonates and infants.

The impairment in visual cortical plasticity was accompanied by a dramatic transcriptional remodeling of the visual cortex. RNA-seq analysis revealed an upregulation of transcripts significantly enriched in pathways related to myelination in ABX-treated mice. In contrast, the transcripts downregulated in the ABX visual cortex clustered in ECM-related annotations. The ECM is a dynamic macromolecular network that undergoes quick changes during neurodevelopment, transitioning from a diffuse, immature structure to a more organized, net-like formation known as PNN ^108,109^. These mature ECM structures emerge at the end of CP, acting as a brake on neuroplasticity in the adult brain ^60,110^. Analysis of PNN architecture in the primary visual cortex demonstrated no changes between ABX and CTRL animals. However, PV+ interneurons displayed a significant increase in their density upon ABX-induced potential dysbiosis. Given the central role of PV+ interneurons in regulating cortical inhibition and CP plasticity, this finding suggests that gut dysbiosis may alter the maturation and functional organization of inhibitory circuits, potentially leading to an imbalance in excitation/inhibition (E/I) ratio. Intriguingly, altered E/I balance is considered a core pathophysiological feature of several neurodevelopmental disorders, including autism spectrum disorders, both in animal models ^111,112^ and human patients ^113,114^. Gut microbiota alterations have also been consistently associated with neurodevelopmental disorders ^9,10,115,116^, further supporting a potential mechanistic link between dysbiosis, inhibitory circuit dysfunction, and altered cortical plasticity.

Our transcriptome analyses unveiled a significant overrepresentation of collagen related terms. Collagen is not a major constituent of PNN ^117^, but rather a component of a more plastic, developmentally active ECM ^118–120^. Given that collagen levels decline in the adult brain ^120^, the prominence of collagen-related annotations in our data suggests that the ECM in CTRL mice may reflect a state of ongoing maturation, which might be compromised in a condition of ABX-induced intestinal microbe disruption. Finally, the enrichment of basal membrane- and collagen- related transcripts prompted us to investigate BBB permeability, which was indeed increased upon ABX treatment. This finding suggests that a disrupted gut microbiota may compromise BBB function, potentially contributing to the observed impairment in experience-dependent plasticity during the CP. This interpretation is consistent with evidence that signals from the intestinal microbiota may affect BBB integrity ^121–124^. In germ-free mice BBB displays a higher permeability associated with decreased expression of occludin and claudin-5 ^125^. Similarly, antibiotic cocktail-induced dysbiosis has been shown to alter tight junction expression in the hippocampus and to cause cognitive deficits in mice ^126^. Although the precise mechanisms linking microbiota alterations to BBB dysfunction remain incompletely understood, several pathways have been proposed, including neural (vagal and sympathetic), immune, and endocrine signaling, as well as the action of microbial metabolites such as SCFA ^127^. In line with this, our analysis revealed a reduction in SCFA-producing taxa in ABX mice. SCFA are known to support BBB integrity ^124,128,129^ and modulate functional neuronal plasticity ^75,130^, thus their reduction may represent a mechanistic link between microbiota alterations, BBB disruption, and impaired OD plasticity during the CP. While the causal chain remains to be experimentally established, our findings raise the possibility that microbiota-dependent regulation of BBB permeability represents a previously underappreciated mechanism controlling CP plasticity.

Successful myelination depends on the differentiation of oligodendrocyte progenitor cells into myelinating oligodendrocytes, a process regulated by both intrinsic signals and environmental influences ^131^. The gut microbiota could play a role in this regulation, although its implication in myelin plasticity and the underlying mechanisms remain incompletely understood ^132^. Emerging evidence indicates that bacterial communities may create very different “biochemical niches”, some permissive and others inhibitory for myelin generation or maintenance ^133,134^. Rodents undergoing early-life ABX exposure display hypermyelination in the prefrontal cortex with respect to their control littermates ^49,135–138^, strengthening the role of the microbiota in modulating myelin assembly in the juvenile brain ^139^. Our transcriptome results are in line with this literature and were partially confirmed by western blot analysis of key myelin-related proteins. LINGO1, an inhibitory regulator of oligodendrocyte precursor cell differentiation ^74^, was reduced in ABX-treated mice. As lower LINGO1 level may indicate increased myelination, this result supports the possibility that ABX-driven dysbiosis during postnatal developmental windows may contribute to altered myelin dynamics in the sensory systems.

Finally, as myelination and inhibitory circuit maturation has been demonstrated to act as a functional brake on visual cortical plasticity during the CP ^59,63,140^, targeting the gut microbiota through ABX administration may impair experience-dependent plasticity by impacting these processes. However, further investigation is needed to confirm this hypothesis.

### Transplanting the critical period gut microbiota restores functional plasticity in adult mice

To causally link the juvenile intestinal microbiota to CP plasticity, we performed a FT experiment in adult, non-plastic mice. Strikingly, the adults receiving the “critical period gut microbiota” exhibited a significant OD shift, confirming that juvenile intestinal microbes are not only necessary but also sufficient to promote cortical plasticity. Importantly, microbial taxa analysis revealed significant differences in the composition of young and adult mice, and demonstrated that young donor “pro-plasticity” bacteria were able to colonize the intestine of adult recipient animals. We finally dissected the microbial signature associated with experience-dependent plasticity by comparing plastic vs non plastic microbiota datasets and revealed eleven bacterial taxa. A subgroup of genera displaying post-transplant increase (i.e., *Ligilactobacillus, Bacteroides, Roseburia, Oscillibacter, [Eubacterium] xylanophilum group, Lachnoclostridium* and *Lachnospiraceae_UCG-006*), are known for their ability to produce SCFA, neuroactive compounds that have been involved in brain plasticity ^75,130,141–143^.

### Translational perspectives

For the first time our work demonstrates that the gut microbiota do not merely influence brain function but may also shape the processes through which neural circuits are stabilized during critical developmental windows, with potentially permanent consequences.

CPs represent developmental time-periods during which experience shapes the “stability landscape” of neural circuits, promoting specific patterns of connectivity that become functionally preferred and unchangeable <u>(Knudsen 2004)</u>. Because plasticity is highly restricted outside these windows, perturbations occurring during the CP can have persistent effects on circuit architecture and function. In this view, our findings identify the gut microbiota as a key endogenous factor capable of influencing the maturation and stabilization of the cortical network during a developmental stage when neural circuits are highly predisposed to remodelling. These results carry important clinical implications for antibiotic use in infants, whose gut microbiota is itself still undergoing maturation and is particularly susceptible to exogenous stimuli. By linking antibiotic-induced dysbiosis to alterations in functional cortical plasticity and transcriptional programs, we provide a framework for understanding how early-life microbial perturbations may influence lifelong brain function and vulnerability to neurodevelopmental disorders ^144–146^. While antibiotics are essential to treat serious infections, our results emphasize the importance of carefully balancing their clinical benefits against potential developmental consequences. Thus, our work could inform future clinical guidelines and protect against potentially deleterious and long-lasting consequences of antibiotic-induced microbial imbalance on the infant brain.

Finally, our discovery represents a significant breakthrough by demonstrating that the juvenile plastic phenotype can be effectively transferred to adult mice through fecal microbiota transplantation. Previous research reported transferring of behavioural and hippocampal plasticity-related features from young to adult/aged rodents ^147–150^. However, this is the first time that using an *in vivo* functional model (i.e., visual cortex), characterized by a defined CP of heightened plasticity ^151^, a study demonstrates that plasticity can be re-activated in a system that typically resists neural network remodeling during adulthood. Remarkably, this effect is achieved through a relatively straightforward strategy, modulating the gut microbiota. Such intervention has vast potential application not only in healthy aging but also in treating neurodegenerative diseases and other neurological disorders. Based on these insights and by modulating a highly modifiable player, i.e. the intestinal microbiota, future therapeutic strategies may significantly improve cognitive resilience, favor functional recovery after brain injury, and ultimately transform clinical care for a range of brain pathologies.

## Resource availability

### Lead contact

Further information and requests for resources should be addressed to the lead contact, Paola Tognini.

### Materials availability

This study did not generate unique reagents.

### Data availability

RNA-seq data generated in this study has been deposited in the NCBI Gene Expression Omnibus (GEO) database, GEO accession number GSE334555.

Any additional information required to reanalyze the data reported in this paper is available from the lead contact upon request.

## Acknowledgments

We thank all the members of Tognini’s team for insightful comments and feedback regarding the experiments. Special thanks to Dr. Maria Grazia Giuliano for her assistance with Western Blots. We also thank Cecilia and Sara Ciampi, Francesca Biondi and Dr. Silvia Burchielli for their help in the mouse facility. This study was in part funded under the National Recovery and Resilience Plan (PNRR), Mission 4 Component 2 Investment 3.1 of the Italian Ministry of University and Research funded by the European Union—NextGenerationEU, “Biorobotics Research and Innovation Engineering Facilities” project (Project identification code IR0000036—CUP J13C22000400007); PNRR YOUNG MSCA_0000081 iNsPIReD, and Italian Ministry of University and Research PRIN 2022-BAGEL, CUP J53C24003080001 to P.T. The work of K.C., S.A., and P.B. was in part supported by NIH grant R01GM123558 to P.B.

## Author Contribution

FD performed all experiments, analyzed the data, and prepared the manuscript figures. KCA and SA conducted the bioinformatic analysis of the RNA-sequencing raw data. AT maintained the mouse colony and helped with ABX administration. SC and MC contributed to stool collection, fecal DNA extraction, and immunofluorescence experiments. PB supervised the bioinformatic analysis and provided feedback on the manuscript. PT conceived the project, supervised the experiments, interpreted the data, and wrote the manuscript.

## Supplemental information titles and legends

**Supplementary Figures:** Supplemental Figures 1-3.

**Suppl. Table 1**: Excel file containing all genes from RNA-sequencing in the visual cortex of P28 mice.

**Suppl. Table 2**: Excel file containing all DAVID annotations from GO, KEGG, and Reactome pathway enrichment analyses performed on differentially expressed genes identified by RNA-sequencing between CTRL and ABX groups, as well as on genes overlapping between those downregulated in ABX mice and the Matrisome dataset.

## MATERIAL AND METHODS

### Animals

All experiments were carried out in accordance with the European Directives (2010/63/EU), and were approved by the Italian Ministry of Health.

All the experiments were performed on female C57BL/6J (JAX #00064), housed (2-3 per cage) in individually ventilated cages to safely maintain the microbiota composition. Weaning was performed on P21–23. Animals were kept in rooms at 22°C with a standard 12 h light–dark cycle. Food (standard diet, 4RF25 GLP Certificate, Mucedola) and water were available *ad libitum* and changed weekly.

### Antibiotic cocktail (ABX) treatment

Antibiotic cocktail (ABX) was administered via autoclaved drinking water and consisted of vancomycin (Duchefa Biochemie Cat# V0155.0005) (0.5 g/L), ampicillin (Sigma-Aldrich Cat#A9518) (1 g/L), and neomycin (Gibco-Life technologies Cat#21810-031) (1 g/L). ABX was refreshed every other day. For intrinsic optical signal (IOS) imaging, RNA sequencing, and Western blot analyses, mice were treated with ABX from weaning until sacrifice (P28–P33), while control littermates received regular drinking water.

### Fecal bacteria DNA extraction and 16S rRNA sequencing

To assess changes in fecal microbiota composition following ABX treatment and fecal microbiota transplantation (FT), fresh fecal pellets were longitudinally collected from the same mice before and after treatment, snap-frozen in liquid nitrogen, and stored at −80°C until processing. Bacterial DNA was extracted using the QIAamp Powerfecal DNA kit (Qiagen, cat # 12830-50), and DNA concentration was quantified by Nanodrop 2000 C Spectrophotometer (ThermoFisher Scientific).

The 16S rRNA sequencing and analysis was performed by Novogene UK Company Ltd.

### Library Preparation and Sequencing

Targeted regions were amplified using specific primers connected with barcodes, followed by selection of PCR products of appropriate size through agarose gel electrophoresis. Equal amounts of PCR products from each sample were pooled, end-repaired, A-tailed, and ligated with Illumina adapters. The prepared libraries were sequenced on a paired-end Illumina platform. The library was checked with Qubit and real-time PCR for quantification, while a bioanalyzer was used for size distribution detection. Quantified libraries were pooled and sequenced on Illumina platforms according to the effective library concentration and data amount required.

### Bioinformatics analysis

Raw sequencing data underwent splicing and filtering to generate clean data. The DADA2 or deblur methods were used to reduce noise and obtain Amplicon Sequence Variants (ASVs). By applying QIIME2’s classify-sklearn algorithm ^152,153^, a pre-trained Naive Bayes classifier is used for species annotation of each ASV. The annotation database of the project is Silva 138.1. Species abundance, alpha diversity and beta diversity were analyzed. Statistical methods, including t-tests, MetagenomeSeq, and LEfSe, were employed to analyze significant differences in species composition between groups. PICRUSt2 was used to predict microbial community functions based on the annotated ASVs.

### Monocular deprivation

Mice were anesthetized with isoflurane (3% induction; 1.5% maintenance) and placed on a heated pad maintained at 37°C. The area surrounding the right eye was cleaned with Povidone-iodine solution using cotton swabs. A thin layer of dexamethasone/tobramycin ointment (Tobradex, Alcon Novartis) was applied to the eye to prevent infection and inflammation. Eyelids were sutured with 2-4 horizontal mattress stitches by using a 6-0 surgical suture. Following surgery, the animals were kept under observation and allowed to recover over the heated pad at 37°C.

### Intrinsic Optical Imaging (IOS)

#### Surgery

Surgery for IOS imaging was performed as described in ^154^. Mice were anesthetized with isoflurane (3% induction; 1.5% maintenance) and head fixed on a stereotaxic frame using ear bars. Body temperature was monitored using a heating pad and a rectal probe to maintain the animals’ body at 37°C. A subcutaneous injection of lidocaine (2%) was provided to anesthetize the local area and the eyes were protected with a dexamethasone-based ointment (Tobradex, Alcon Novartis). The scalp was removed and the skull cleaned with saline. The skin was secured to the skull using cyanoacrylate and a thin layer of cyanoacrylate was poured over the exposed skull to attach a custom-made metal ring (9 mm internal diameter) centered over the binocular visual cortex. A thin layer of clear nail polish was applied over the area to ameliorate optical access. After surgery, the animals were allowed to fully recover inside a heated box and monitored to ensure the absence of any sign of discomfort. Before any other experimental procedure, mice were left to recover for at least 4 days.

#### Imaging and data analysis

Mice were anesthetized with isoflurane (3% induction; 1.5% maintenance) and chlorprothixene anesthesia (1.5 mg/kg, i.p.). Images were visualized using a custom Leica microscope (Leica Microsystems). Red light illumination was provided by 6 individually addressable LEDs (WS2812) attached to the objective (Leica Z6 APO coupled with a Leica PlanApo 2.0X 10447178) by a custom 3D-printed conical holder. Visual stimuli were generated using Matlab Psychtoolbox and presented on a gamma-corrected 24” monitor (C24F390FHU).

Horizontal sine-wave gratings were presented in the binocular portion of the visual field enclosed in a Gaussian envelope spanning -10 to +10 degrees of azimuth and -5 to +60 (full monitor height) degrees of altitude, with a spatial frequency of 0.03 cycles per degree, mean luminance 20 cd/m^2^ and a contrast of 90%. The stimulus consisted of the abrupt contrast reversal of a grating with a temporal frequency of 4 Hz for 1 second, time-locked with a 12-bit depth acquisition camera (PCO edge 5.5) using a parallel port trigger. The interstimulus time was 13 seconds. Frames were acquired at 30 fps with a resolution of 540 x 640 pixels. The signal was averaged for at least 6 groups of 20 trials, stimulating each eye alternatively to prevent biases due to different time in anesthesia for the contralateral and ipsilateral eyes. The signal was then downsampled in time to 10 fps and in space to 270 x 320 pixels. Fluctuations of reflectance (R) for each pixel were computed as the normalized difference from the average baseline (ΔR/R). For each recording, an image representing the mean evoked response was computed by averaging frames between 0.5 to 2.5 seconds after stimulation. The mean image was then low-pass filtered with a 2D average square spatial filter (7 pixels). To select the binocular portion of the primary visual cortex for further analysis, a region of interest (ROI) was automatically calculated on the mean image of the response by selecting the pixels in the lowest 20% ΔR/R of the range between the maximal and minimal intensity pixel ^155^. To weaken background fluctuations a manually selected polygonal region of reference (ROR) was subtracted. The ROR was placed where no clear response, blood vessel artifact or irregularities of the skull were observed ^156^. Mean evoked responses were quantitatively estimated as the average intensity inside the ROI. Ocular dominance was determined through the Ocular Dominance Index (ODI), calculated as 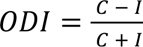, where C and I are the mean response amplitude evoked from contralateral and ipsilateral eye stimulation respectively. Shift in the ocular dominance was evaluated comparing ODI before and after 3 days of MD.

### RNA extraction

Visual cortex samples were homogenized in phenol/guanidine-based QIAzol Lysis Reagent (Qiagen Cat. #79,306). Chloroform was added and the samples were vigorously shaken for 15 s. The samples were left at 20–24 °C for 3 min and then centrifuged (12,000×*g*, 20 min, 4 °C). The upper phase aqueous solution, containing RNA, was collected in a fresh tube and the RNA was precipitated by the addition of isopropanol. Samples were mixed by vortexing, left at room temperature (RT) for 10 min and then centrifuged (12,000×*g*, 15 min, 4 °C). The supernatant was discarded and the RNA pellet was washed in 75% ethanol by centrifugation (7500×*g*, 5 min, 4 °C). The supernatant was discarded and the pellet was left to dry for a minimum of 15 min before resuspending in RNAse free water. RNA concentration was determined by Nanodrop Spectrophotometer (Thermoscientific 2000 C).

### RNA-sequencing

N = 4 biological replicates per experimental group were used in the RNA-seq experiment. Total RNA was extracted as described above. RNA sequencing was performed by Novogene UK Company Ltd.

#### Library preparation and sequencing

For the library preparation, messenger RNA was purified from total RNA using poly-T oligo-attached magnetic beads. After fragmentation, the first strand cDNA was synthesized using random hexamer primers followed by the second strand cDNA synthesis. The library was ready after end repair, A-tailing, adapter ligation, size selection, amplification, and purification. mRNA adapter sequences:

5’ Adapter: 5’-AGATCGGAAGAGCGTCGTGTAGGGAAAGAGTGTAGATCTCGGTGGTCGCCGTATC ATT-3’

3’ Adapter: 5’-GATCGGAAGAGCACACGTCTGAACTCCAGTCACGGATGACTATCTCGTATGCCGT CTTCTGCTTG-3’

The library was checked with Qubit and real-time PCR for quantification and bioanalyzer for size distribution detection.

Quantified libraries were pooled and sequenced on Illumina NovaSeq platforms, according to effective library concentration and data amount.

#### Bioinformatics analysis

The FastQ files were processed through the standard Tuxedo protocol, using Tophat and Cufflinks ^157,158^. Tophat was used to align the RNA-seq reads to the reference genome assembly mm10 and Cufflinks was used to calculate gene expression levels. This protocol outputs the FPKM values for each gene of each replicate. The differential analysis of the FPKM values across all experiments and control groups was conducted with Cyber-T, a differential analysis program using a Bayesian-regularized *t*-test ^159,160^. The *p*-value threshold used for determining differential expression was 0.05.

Gene ontology (GO), Kyoto Encyclopedia of Genes and Genomes (KEGG) and Reactome enrichment pathway analysis was performed using the functional annotation chart of DAVID tool, as it allows the integration of multiple databases in one output ^161,162^.

### Immunofluorescence staining

Immunofluorescence (IF) staining of PV^+^ interneurons and WFA^+^ cell-ensheathing PNN was performed on brain slices of ABX-treated and CTRL mice at P30.

Briefly, mice were anesthetized with chloral hydrate (20ml/Kg BW) and perfused via intracardiac infusion, first with PBS and then with 4% paraformaldehyde (PFA, w/vol, dissolved in 0.1 M phosphate buffer, pH 7.4). The brain was quickly removed and post-fixed overnight in PFA at 4 °C, then transferred to a 30% sucrose (w/vol) solution until usage. Coronal brain slices (45 µm) were sectioned with a dry ice-cooled microtome and free-floating collected. Four sections containing the V1 region were selected for every mouse and used for IF staining. Immunodetection was performed using indirect antigen labeling with prior blocking of nonspecific binding sites through bovine serum albumin (BSA) buffers to minimize background signal.

For PNN detection, sections were incubated for 1 hour at RT in a blocking solution composed of 3% BSA (w/vol) in PBS, followed by an overnight incubation at 4°C with the *N*-acetylgalactosamine-binding *Wisteria floribunda* agglutinin (WFA) (Cat# B-1355) diluted 1:200 in the same solution used for blocking. The following day, sections underwent 3 washing of 10 min/each in PBS, and then were incubated for 2 hours at RT with Streptavidin 555 (Invitrogen Cat# S21381) diluted 1:400 in a solution composed of 3% BSA in PBS.

To label PV+ neurons, sections underwent additional blocking with 10% BSA (w/vol) and 0.3% Triton X-100 (vol/vol) in PBS, and then incubated overnight at 4°C with anti-PV monoclonal primary antibody (Synaptic System Cat# 195308) diluted 1:1000 in a solution of PBS with 1% BSA (w/vol), and 0.1% Triton X-100 (vol/vol). The following day, sections were washed with PBS and incubated for 2 hours at RT with Alexa Fluor 488 anti-guinea pig secondary antibody (Invitrogen Cat# A11073) diluted 1:500 in a solution of PBS with 1% BSA (w/vol) and 0.1% Triton X-100 (vol/vol). Finally, after 3 PBS washing, sections were mounted using Hoechst 405 staining solution.

Imaging was performed on a Leica STELLARIS 5 confocal microscope (LeicaMicrosystem, Wetzlar, Germany) using a dry 20x objective.

Images (512 × 512 pixels) were acquired under standardized conditions (fixed light intensity, digital gain, and exposure time) and exported in .lif format. Region of interest (ROI) sections, corresponding to the V1 regions, were defined based on the Mouse Brain Atlas (Paxinos and Franklin’s the Mouse Brain in Stereotaxic Coordinates).

PV^+^ cells and WFA^+^ PNN countings were performed using a custom MATLAB-based graphical user interface called CounTastic and available at Zenodo ^163^. Density was quantified dividing the number of identified objects (PV^+^ cells and WFA^+^ PNN) per unit of surface area (mm^2^). Staining intensity of each individual PNN and PV cell was quantified by averaging the values of the pixels belonging to the object, segmented from a small (40 × 40 pixels) patch centered on its (x,y) coordinates.

### Protein extraction and Western Blot

For total protein extracts, visual cortex samples were harvested, snapped frozen in liquid nitrogen and stored at −80 °C until usage. Samples were homogenized in modified RIPA buffer (50 mM Tris pH8, 150 mM NaCl, 5 mm EDTA, 15 mM MgCl2, 1% NP40) plus protease inhibitors (DOC 10%, PIC 20X, PMSF 100 mM). Samples were sonicated for 1 min and 10 sec on ice (10 sec on/10 sec off) and centrifuged for 15 min at 14.000×g, at 4°C. The supernatant was recovered and the protein concentration was determined by Bradford assay (Biorad, Cat# 5000006) using a Nanodrop Spectrophotometer (Thermoscientific 2000 C).

8% SDS-PAGE was performed to check Lingo 1 levels. 12% SDS-PAGE was performed to analyze Olig2 and MBP proteins. Samples were blotted onto nitrocellulose membranes (Biorad, Cat# 1620112) and blocked in 5% milk+0.2% Tween in Tris-buffered saline (TBS) for 1 h at RT. The nitrocellulose membrane was incubated at 4 °C overnight with the following antibodies: rabbit polyclonal antibody Anti-Lingo1 (Merck Millipore, Cat. # 07-678), rabbit polyclonal antibody Anti-Olig2 (Merck Millipore, Cat. # AB9610), mouse monoclonal antibody Anti-MBP (BioLegend, Cat. # 836504). Blots were then washed 3 times in TTBS for 30 min, incubated with mouse anti-rabbit IgG-HRP (Santa Cruz Biotechnology, Cat. # sc-2357) or goat anti-mouse IgG-HRP secondary antibodies diluted (1:1000) in 2.5% milk+0.1% Tween in TBS for 1 h at RT. The membranes were then rinsed three times in TTBS and incubated in enhanced chemiluminescent substrate (Biorad Clarity Western ECL substrate, Cat# 170 5060, and Life Technologies Supersignal West Femto Maximum Sensitivity Substrate, Cat# 34095) and acquired through a Chemidoc XRS instrument. Bands densitometry was analyzed through the ImageJ software.

### Evan’s Blue extravasation assay

Blood–brain barrier (BBB) permeability was assessed by Evans Blue (EB) extravasation assay in mice aged P35–P38. EB dye (Merck Sigma-Aldrich, Cat# E2129) (1% w/v) was prepared in sterile saline and administered intraperitoneally at a dose of 8 mL/kg body weight. After injection, mice were returned to their cages and maintained in a warm environment for 2 h to allow systemic circulation of the dye. Mice were then anesthetized with chloral hydrate according to body weight and transcardially perfused with PBS 1X to remove intravascular dye. Brains were then rapidly dissected, snap-frozen in liquid nitrogen, and stored at −80 °C until usage.

For EB quantification, half brains were weighed using an analytical balance and homogenized in 750 μL of 50% trichloroacetic acid (CARLO ERBA Reagents srl, Cat# 411525) (TCA, w/v) using scissors and a tissue grinder. The remaining halves were subsequently processed as technical replicates. Homogenates were centrifuged at 10,000 × g for 20 min at RT, and the resulting supernatant was collected and diluted 1:1 with ethanol. EB absorbance was measured at λ=620 nm using a NanoDrop 2000c spectrophotometer (Thermo Fisher Scientific). Quantification was performed using external EB standards (25–1000 ng/mL) prepared in 50% TCA/ethanol (1:1). EB concentration was determined from a linear standard curve and expressed as nanograms of EB per milligram of brain tissue.

### Fecal transplantation

Adult female C57BL/6J recipient mice (P120), housed 2–3 per cage after weaning (P21), were randomly assigned to one of three experimental groups: (i) fecal microbiota transplantation (FT) from juvenile female C57BL/6J donors (P28), (ii) FT from adult female conspecific donors, or (iii) sterile PBS inoculation as vehicle control. Prior to transplantation, recipient mice were treated with ABX (vancomycin (0.5 g/L), ampicillin (1 g/L), neomycin (1 g/L) and metronidazole (Merck Sigma-Aldrick Cat# M3761) (1 g/L)) in the drinking water for 1 week. Following a 2-day washout period, FT was administered by oral gavage once a day for five total administrations, following previously described procedures ^75,115,164^. To minimize stress, gavage was performed over three consecutive days, followed by a 2-day rest period, and then resumed for two additional days. For FT, freshly collected donor fecal pellets were suspended in sterile PBS, vortexed for 10 min to ensure homogeneous mixing, and filtered through a cell strainer to remove large debris. The resulting suspension was immediately used for transplantation, with each recipient receiving 200 µL by oral gavage. After FT, mice from each experimental group were housed in separate autoclaved cages for four weeks to allow microbiota engraftment. Fecal pellets from recipient mice were collected before ABX treatment and four weeks after FT for subsequent analyses, see the section ‘‘Fecal bacterial DNA extraction and 16S rRNA sequencing’’.

### Statistical Analysis

The sample sizes were based on prior studies and are indicated in the figure legend for each panel. Whenever possible, quantification and analyses were performed blind to the experimental condition. The majority of statistical analyses were performed using GraphPad Prism version 7 (GraphPad Software, San Diego, CA, USA). All data are represented as the mean ± SEM unless otherwise stated. N’s and circles over the bar plots represent single animals unless otherwise stated. Statistical significance was defined in the figure panels as follows: *P ≤ 0.05, **P ≤ 0.01 and ***P ≤ 0.001.

#### Assessment of the ABX treatment

Differences in fecal bacterial DNA concentration in stool (ng/mg) of CTRL and ABX mice were tested for significance using unpaired nonparametric Mann-Whitney U test, as normality and homoscedasticity assumptions were not respected. Differences in the absolute body weight (g) between CTRL and ABX at P21 and P28 were tested using RM Two-way ANOVA, followed by Sidak’s post hoc multiple comparisons test. Differences on the percentage of body weight from baseline between CTRL and ABX were tested for significance using Mann-Whitney U test ****p<0.0001.

#### Gut microbiota analysis

To test whether two or more groups of samples were significantly different, ANOSIM and principal coordinate analysis (PCoA) were calculated using the Python library scikit-bio (http://scikit-bio.org/) with, respectively, the skbio.stats.distance.anosim and skbio.stats.ordination.pcoa functions. Alpha diversity significance was calculated using the Mann-Whitney test.

#### IOS experiments

Variations of ODI after 3 days of MD were tested for significance within each group with paired samples t-test, as normality and homoscedasticity assumptions were respected.

#### RNA sequencing analysis

Transcripts from the RNAseq with a p-value<0.05 were used to perform gene ontology analysis.

#### Immunofluorescence analysis

Differences in cumulative probability of staining intensity in cells were tested for significance with Kolmogorov-Smirnov test. Differences in staining intensity (averaged for each animal) between CTRL and ABX mice were tested for significance with unpaired parametric t-test. The same test was used for the analysis of PV^+^ cells and WFA^+^ PNN density.

#### Western blot analysis

Differences in the relative band intensity between CTRL and ABX mice were tested for statistical significance using unpaired parametric t-test when normality and homoscedasticity assumptions were respected ((i) MBP and ii) Lingo-1), or using Mann-Whitney U test when the samples did not follow a normal distribution ((iii) Olig-2).

