## Supplementary material for "The Critical Period Microbiota Shape Brain Plasticity": Suppl. Figures

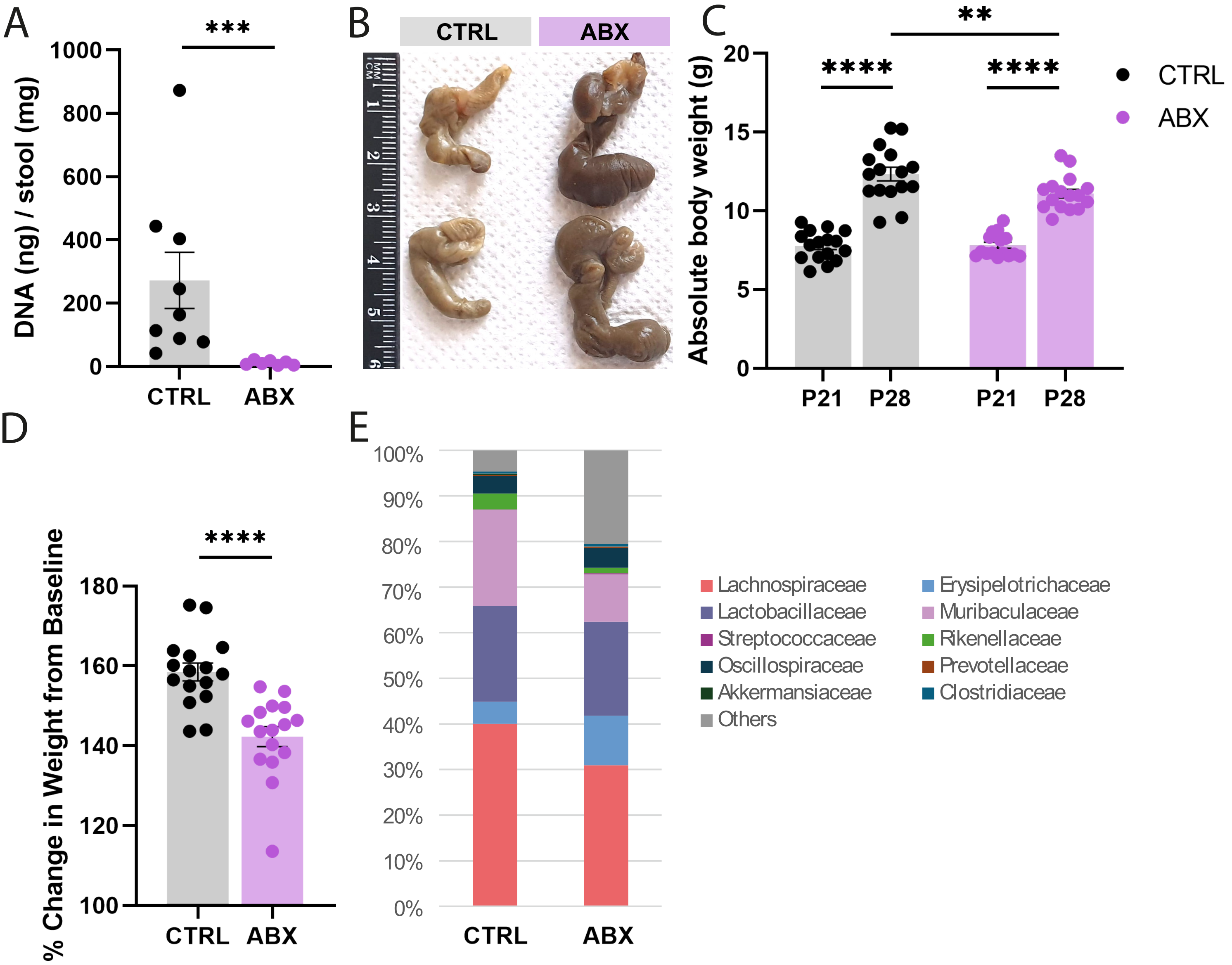


**SUPPL. FIGURE 1.** Assessment of ABX treatment efficacy. **(A)** Fecal bacterial DNA concentration in CTRL and ABX mice after one week of treatment (P28). N=8 animals/group. Mann-Whitney U test ***p=0.0002. **(B)** Representative images comparing cecum size between CTRL (n=2) and ABX (n=2) mice. **(C)** Absolute body weight (g) of CTRL and ABX mice at P21 (weaning) and P28 (after one week of ABX treatment). Body weight measurements N=16 animals/group. RM Two-way ANOVA time*treatment interaction p=0.034, main effect of time p<0.0001, main effect of treatment p<0.0001, multiple comparisons Sidak post-hoc test, P21 CTRL versus P28 CTRL ****p<0.0001, P21 ABX versus P28 ABX ****p<0.0001, P28 CTRL versus P28 ABX **p=0.008. **(D)** Percentage of body weight gain from baseline (P21) to P28. Mann-Whitney U test ****p<0.0001. **(E)** Relative abundance of the ten most prevalent bacterial families in CTRL and ABX mice. Less represented bacterial families are grouped under "Others." N=6 animals/group.

Error bars represent SEM. Circles represent single experimental subjects.

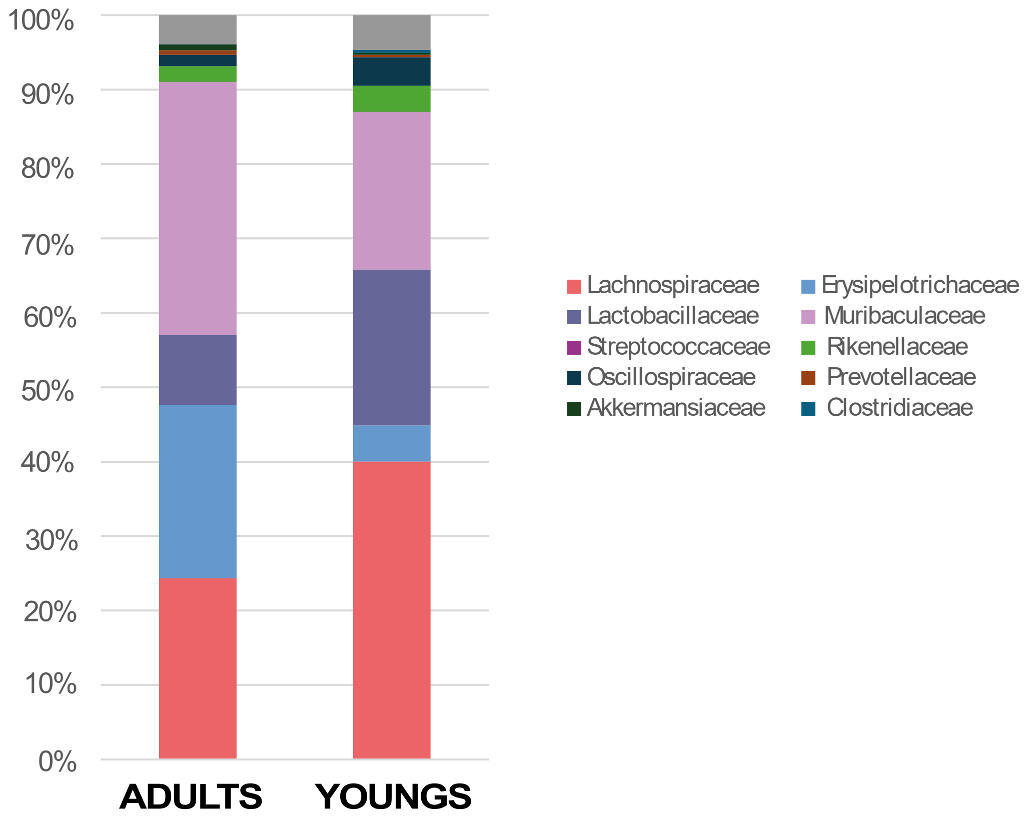


**SUPPL. FIGURE 2.** Relative abundance of the ten most prevalent bacterial families in adult and young mice. Less represented bacterial families are grouped under "Others." N=6 animals/group.

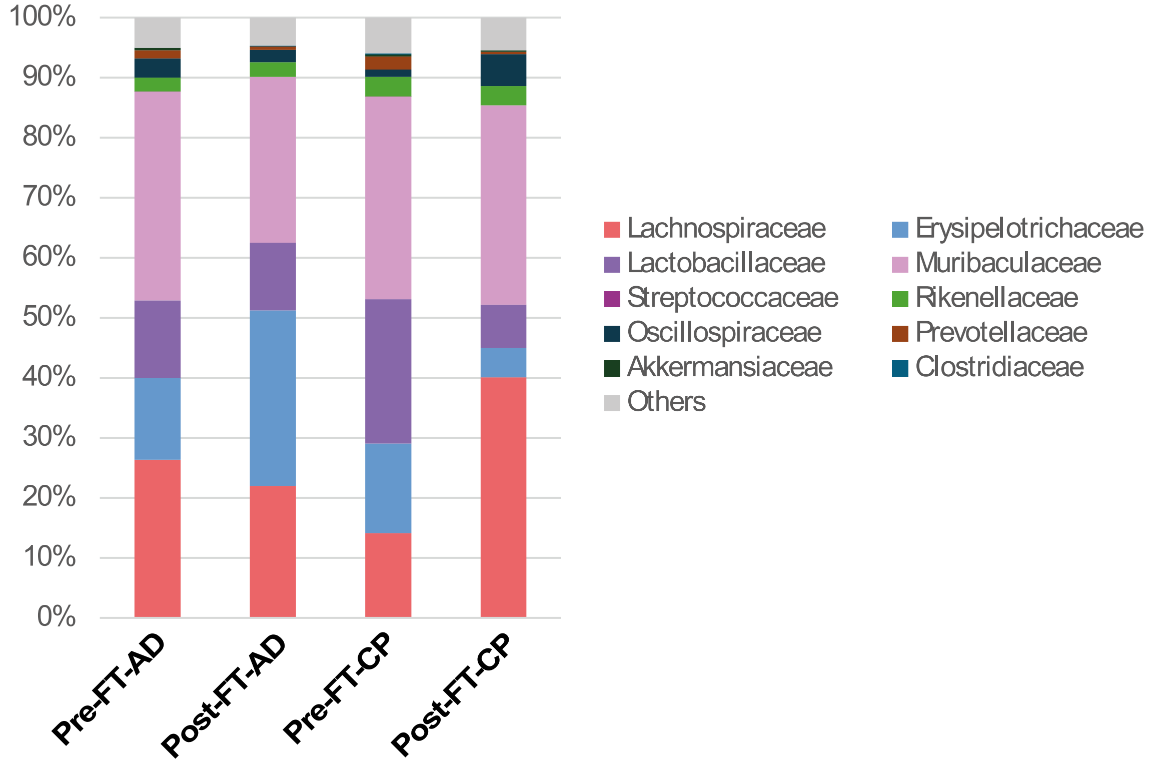


**SUPPL. FIGURE 3.** Relative abundance of the ten most prevalent bacterial families in fecal transplant recipient mice. Less represented bacterial families are grouped under "Others". Pre-FT-AD= pre-FT from adult donors; Post-FT-AD= post-FT from adult donors; Pre-FT-CP= pre-FT from young donors; Post-FT-CP= post-FT from young donors. N=6 animals/group.
